# Metagenomic sequencing suggests a diversity of RNA interference-like responses to viruses across multicellular eukaryotes

**DOI:** 10.1101/166488

**Authors:** Fergal M. Waldron, Graham N. Stone, Darren J. Obbard

## Abstract

RNA interference (RNAi)-related pathways target viruses and transposable element (TE) transcripts in plants, fungi, and ecdysozoans (nematodes and arthropods), giving protection against infection and transmission. In each case, this produces abundant TE and virus-derived 20-30nt small RNAs, which provide a characteristic signature of RNAi-mediated defence. The broad phylogenetic distribution of the Argonaute and Dicer-family genes that mediate these pathways suggests that defensive RNAi is ancient and probably shared by most animal (metazoan) phyla. Indeed, while vertebrates had been thought an exception, it has recently been argued that mammals also possess an antiviral RNAi pathway, although its immunological relevance is currently uncertain and the viral small RNAs are not detectably under natural conditions. Here we use a metagenomic approach to test for the presence of virus-derived small RNAs in five divergent animal phyla (Porifera, Cnidaria, Echinodermata, Mollusca, and Annelida), and in a brown alga—which represents an independent origin of multicellularity from plants, fungi, and animals. We use metagenomic RNA sequencing to identify around 80 virus-like contigs in these lineages, and small RNA sequencing to identify small RNAs derived from those viruses. Contrary to our expectations, we were unable to identify canonical (i.e. *Drosophila-*, nematode- or plant-like) viral small RNAs in any of these organisms, despite the widespread presence of abundant micro-RNAs, and transposon-derived somatic Piwi-interacting piRNAs in the animals. Instead, we identified a distinctive group of virus-derived small RNAs in the mollusc, which have a piRNA-like length distribution but lack key signatures of piRNA biogenesis, and a group of 21U virus-derived small RNAs in the brown alga. We also identified primary piRNAs derived from putatively endogenous copies of DNA viruses in the cnidarian and the echinoderm, and an endogenous RNA virus in the mollusc. The absence of canonical virus-derived small RNAs from our samples may suggest that the majority of animal phyla lack an antiviral RNAi response. Alternatively, these phyla could possess an antiviral RNAi response resembling that reported for vertebrates, which is not detectable through simple metagenomic sequencing of wild-type individuals. In either case, our findings suggest that the current antiviral RNAi responses of arthropods and nematodes are highly diverged from the ancestral metazoan state, and that antiviral RNAi may even have evolved independently on multiple occasions.

**Author summary:** The presence of abundant virus-derived small RNAs in infected plants, fungi, nematodes, and arthropods suggests that Dicer-dependent antiviral RNAi is an ancient and conserved defence. Using metagenomic sequencing from wild-caught organisms we show that antiviral RNAi is highly variable across animals. We identify a distinctive group of virus-derived small RNAs in a mollusc, which have a piRNA-like length distribution but lack key signatures of piRNA biogenesis. We also report a group of 21U virus-derived small RNAs in a brown alga, which represents an origin of multicellularity separate from that of plants, fungi, and animals. The absence of virus-derived small RNAs from our samples may suggest that the majority of animal phyla lack an antiviral RNAi response or that these phyla could possess an antiviral RNAi response resembling that reported for vertebrates, which is not detectable through simple metagenomic sequencing of wild-type individuals. In addition, we report abundant somatic piRNAs across anciently divergent animals suggesting that this is the ancestral state in Bilateria. Our study challenges the widely-held assumption that most invertebrates possess an antiviral RNAi pathway likely similar to that seen in *Drosophila*, other arthropods, and nematodes.

## Introduction

RNA interference-related (RNAi) pathways provide an important line of defence against parasitic nucleic acids in plants, fungi, and most animals (1–5). In plants and fungi, which lack a distinct germline, Dicer and Argonaute-dependent RNAi responses suppress the expression and replication of viruses and transposable elements (TEs) through a combination of target cleavage and/or heterochromatin induction (6,7). This gives rise to a characteristic signature of short interfering RNAs (siRNAs) derived from both TEs and viruses (8–12). In contrast, the best-studied animal (metazoan) lineages display two distinct signatures of defensive RNAi. First, reminiscent of plants and fungi, arthropods and nematodes exhibit a highly active Dicer-dependent antiviral pathway that is characterised by copious virus-derived siRNAs (viRNAs) peaking sharply between 20nt (e.g. Lepidoptera) and 22nt (e.g. Hymenoptera). These are cleaved from double-stranded viral RNA by Dicer, and loaded into an Argonaute-containing complex that targets virus genomes and transcripts via sequence complementarity (13,14). Second, and in contrast to plants and fungi, animals also possess a Piwi-dependent (piRNA) pathway that provides a defence against TEs in germline (*Drosophila* and mammals) and/or somatic cells (e.g. 15–17). This pathway is usually characterised by a broad peak of 26-30nt small RNAs bound by Piwi-family Argonaute proteins, and comprises both 5’U primary piRNAs cleaved by homologs of *Drosophila* Zucchini from long ‘piRNA cluster’ transcripts (see 18), and secondary piRNAs generated by ‘Ping-Pong’ amplification. In most animals it is thought to target TE transcripts for cleavage, and genomic copies for heterochromatin induction (19).

The presence of abundant viRNAs in infected plants, fungi, nematodes, and arthropods suggests that Dicer-dependent antiviral RNAi is an ancient and conserved defence (1,2). However, RNAi has been entirely lost in some lineages such as *Plasmodium* (20), some trypanosomes (21), and some *Saccharomyces* (22), and/or extensively modified in others. For example, antiviral RNAi was long thought to be absent from vertebrates (23,24), at least in part because their viRNAs cannot easily be detect by high-throughput sequencing of the total small-RNA pool from wild-type individuals (23–28). Recently, it has been suggested that vertebrates also possess a functional virus-targeting RNAi pathway in tissues lacking an interferon response (29–31) and/or in the absence of viral suppressor of RNAi (30,32,33). However, there is still debate as to whether this occurs under natural conditions, and whether or not it represents an immunologically relevant defence (compare 34,35).

Despite this clear interest in the phylogenetic distribution of antiviral RNAi, comprehensive experimental studies of antiviral RNAi in animals are not available, with studies instead focussing on arthropods such as insects (reviewed in 36,37), crustaceans (38, and reviewed in 39), chelicerates (40)), and on nematodes (41–43) and vertebrates (23–25,27,29–33,44). In particular, there have been few attempts to identify viRNAs in ‘early-branching’ animal lineages such as Porifera or Cnidaria, in divergent Deuterostome lineages such as Echinodermata or Urochordata, or in Lophotrochozoa (including the large phyla Annelida and Mollusca; See Fig 1 for the known distribution of RNAi-pathways across the Metazoa).

**Fig 1.**
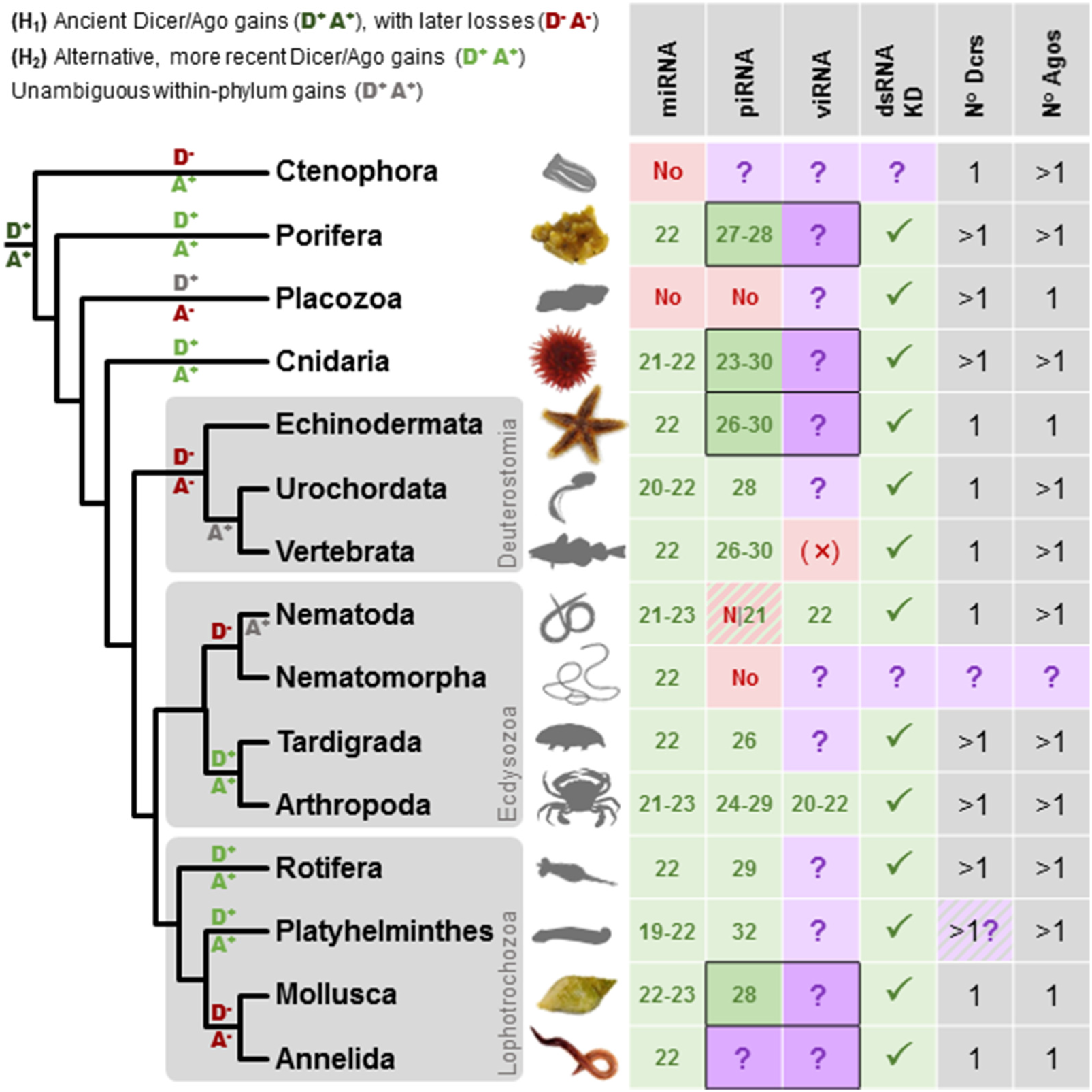
Distribution of small RNA pathways across the Metazoa. Phylogeny of selected metazoan (animal) phyla (topology follows 172) with a table recording the reported range of modal lengths for miRNAs, piRNAs, and viRNAs detectable by bulk sequencing from wild-type organisms (miRNA modes taken from miRbase). Entries marked ‘No’ have been reported to be absent, and those marked ‘?’ are untested. Focal taxa in this study are marked in colour, and the target table entries are outlined. Vertebrate viRNAs are marked ‘(×)’ as mammalian virus-derived small RNAs are only detectable in tissues and experimental systems lacking viral suppressors of RNAi and/or an interferon response (29–33). Note that piRNAs are absent from some, but not all, nematodes (55). The column ‘dsRNA KD’ records whether dsRNA knockdown of gene expression using long dsRNA (i.e. a Dicer substrate) has been reported, as this may suggest the presence of an RNAi pathway capable of producing viRNAs from replicating viruses. The ‘Dcrs’ and ‘Agos’ columns record the inferred number of Dicers and (non-Piwi) Argonautes ancestrally present in each phylum, although the number of Dicers in Platyhelminthes is contentious as the putative second Dicer lacks the majority of expected Dicer domains. Broadly speaking, there are two competing hypotheses for the histories of Dicers and (non-Piwi) Argonautes in animals, (e.g. 46,173,174). The first (H_1_), posits that an early duplication in Dicer and/or Argonaute (marked D^+^ and A^+^ in dark green on the phylogeny) gave rise to at least two very divergent homologues of each gene in the lineage leading to the metazoa, followed by subsequent losses (D^−^ and A^−^ in dark red). The second (H_2_), suggests that divergent homologues are the result of more recent duplications (D^+^ and A^+^ in pale green), and where homologs have high divergence it is as a result of rapid evolution. Note that these hypotheses are independent for Argonautes and Dicers, and one may be ancient but the other recent. For Dicers, at least, the ‘ancient’ duplication is arguably better supported (46), although it remains extremely difficult to determine orthology between the duplicates. In addition, Dicers and Argonautes have unambiguously diversified within some phyla (important examples marked A^+^ and D^+^ in grey)—as seen for the large nematode-specific WAGO clade of Argonautes (reviewed in 133), and the multiple Argonautes in vertebrates.

Broadly consistent with a wide distribution of antiviral RNAi, Argonaute and Dicer genes are detectable in most animal genomes (Fig 1; 45–48). However, while Dicer and Argonaute genes would be necessary for a canonical antiviral RNAi response, their presence is insufficient to demonstrate one, for two reasons. First, these genes also have non-defensive roles such as transcription regulation through miRNAs (see 49,50)—and a single gene can fulfil multiple roles. For example, whereas in *Drosophila* there is a distinction between the Dcr2-Ago2 antiviral pathway and the Ago1-Dcr1 miRNA pathway (e.g. 51), in *C. elegans* a single Dicer is required for the biogenesis of both miRNAs and viRNAs (Fig 1; 41,52,53). Second, RNAi pathways are labile over evolutionary timescales, with regular gene duplication, loss, and change of function (e.g. 54–56). For example, the Piwi-family Argonaute genes that mediate anti-TE defence in Metazoa were ancestrally present in eukaryotes, but were lost independently in plants, fungi, brown algae, most nematodes, and dust mites (2,46,55–57). In contrast, non-Piwi Argonautes were lost in many alveolates, excavates and Amoebozoa (57,58) while Piwi genes were retained in these lineages. At the same time, new RNAi mechanisms have arisen, such as the 22G RNAs of nematodes (55,59,60) and the recent gain of an antiviral role for Piwi in *Aedes* mosquitoes (61,62). Taken together, the potential for multiple functions, and for gains and losses of function, make it challenging to confidently predict the phylogenetic distribution of antiviral RNAi from the distribution of the required genes alone (see 48).

Thus, although antiviral RNAi is predicted to be shared by most extant eukaryotes (see 63,64), in the absence of experimental studies, its distribution across animal phyla remains largely unknown (Fig 1). This contrasts sharply with our knowledge of other RNAi-related pathways, such as the micro-RNA (miRNA) mediated control of gene expression, which is conserved across plants, brown algae, fungi, and almost all animals (65), and the presence of TE-derived piRNAs in most animals: Porifera (15,66), Cnidaria (15,67), Ctenophora (68), Vertebrata (69,70), Arthropoda (71–74), some Nematoda (55, but see 75), Platyhelminthes (76), but not Placozoa (15). In eukaryotes that lack direct experimental evidence for viRNAs, the presence of an inducible RNAi response to experimentally applied long double-stranded RNA might indicate a potential for antiviral RNAi (Fig 1). This has been reported for Excavata (77), Heterkonta (78) Amoebozoa (79), trypanosomes (80), and among Metazoa in Porifera (81), Cnidaria (82), Placozoa (83), Arthropoda (84), Nematoda (85), and several lineages of Lophotrochozoa including planarian flatworms (86), bivalve molluscs (87), rotifers (88) and annelids (89).

Thus, although circumstantial evidence suggests a near-universal potential for antiviral RNAi in animals, we still lack experimental evidence of exogenous viral processing. This knowledge gap is probably attributable, in part at least, to the challenges associated with isolating and culturing non-model animals and their natural viral pathogens in the lab. Here we seek to examine the phylogenetic distribution of viRNAs, and thus elucidate the phylogenetic distribution of a canonical (i.e. *Drosophila-*, nematode- or plant-like) antiviral RNAi response, through metagenomic sequencing. We combine rRNA-depleted RNA sequencing with small-RNA sequencing to detect both viruses and viRNAs in pooled samples of six deeply divergent lineages. First, we include two early branching metazoan phyla: a sponge (*Halichondria panicea*: Porifera, Demospongiae) and a sea anemone (*Actinia equina*: Cnidaria, Anthozoa) that branch basally to the divergence between deuterostomes and protostomes (Fig 1). Second, a starfish (*Asterias rubens*: Echinodermata, Asteroidea) that branches basally to vertebrates within the Deuterostomia. Third, two divergent species of Lophotrochozoa—the clade which forms the sister group to Ecdysozoa within the protostomes: a dog whelk (*Nucella lapillus*: Mollusca, Gastropoda) and earthworms (Annelida, Oligochaeta). Finally, to explore the deep history of antiviral RNAi within the eukaryotes, we included the brown alga *Fucus serratus* (Phaeophyceae, Heterokonta), which represents an origin of multicellularity separate from that of plants, fungi, and animals.

This metagenomic approach circumvents the need to isolate and/or culture non-model organisms in the laboratory, and can capitalise on the high diversity of viruses naturally infecting individuals in the wild. It also avoids any artefactual that might result from non-native host-virus combinations, or non-natural infection routes. Surprisingly, although we find viral RNA sequences to be common and sometimes highly abundant, we do not find abundant viRNAs from RNA viruses in most of the sampled species, suggesting that they lack a canonical (i.e. *Drosophila-*, nematode- or plant-like) antiviral RNAi response. Specifically, we detect no viRNAs from RNA viruses infecting the earthworms, the sponge, or the sea anemone, suggesting that insect- or nematode-like antiviral RNAi is absent from these lineages. In contrast, we do detect viRNAs from RNA viruses in the dog whelk and the brown alga. In both cases these viRNAs derive from both strands of the virus. However, in the dog whelk they peak broadly at 26-30nt—as would be expected of piRNAs, but lacking the 5’U or ‘ping-pong’ signature— and in the brown alga they peak sharply at 21nt and are exclusively 5’U. This suggests the presence of distinct antiviral RNAi responses in these two lineages. Finally, we identify primary piRNA-like 26- 30nt 5’U small-RNAs derived from putatively endogenous copies of viruses in the sponge, the starfish, and the dog whelk, and somatic TE-derived piRNAs in all the animal lineages examined, suggesting an origin of somatic piRNAs at least as old as ancestral Bilateria. Taken together, these findings imply that the true diversity of defensive RNAi strategies employed by eukaryotes may have been underestimated, and that antiviral RNAi is either lacking from many animal phyla, or perhaps resembles the RNAi response reported for mammals.

## Results

### New virus-like sequences identified by metagenomic sequencing

Using the Illumina platform, we generated strand-specific 150 nt paired-end sequence reads from ribosome-depleted RNA extracted from metagenomic pools of each of six different species: the breadcrumb sponge (*Halichondria panacea*, Porifera); the beadlet sea anemone (*Actinia equina*, Cnidaria); the common starfish (*Asterias rubens*, Echinodermata); the dog whelk (*Nucella lapillus*, Mollusca); mixed earthworm species (*Amynthas* and *Lumbricus* spp., Annelida), and a brown alga (the ‘serrated wrack’, *Fucus serratus*, Fucales, Phaeophyceae, Heterokonta). See S1 Table for collection data. Gut contents were excluded by dissection, and contaminating nematodes excluded by a PCR screen prior to pooling (Materials and Methods; S1 Table). Reads were assembled separately for each species using Trinity v2.2.0 (90,91), resulting in between 104,000 contigs for the sponge and 235,000 contigs for the earthworms. Metagenomic analysis using Diamond v0.7.11.60 (92) and MEGAN6 (93) suggests the vast majority of these contigs derive for the intended host organism (S1 Fig). Unannotated contigs are provided in supporting file S1 Data. To identify viruses, we used Diamond to search with translated open reading frames (ORFs) from our contigs against all virus proteins from the NCBI nr database, all predicted proteins from Repbase (94), and all proteins from the NCBI RefSeq_protein database (see Materials and Methods). After excluding some low-quality matches to large DNA viruses and matches to phage, this identified nearly 900 potentially virus-like contigs (S2 Data). These matches were examined and manually curated to generate 85 high-confidence virus-like contigs between 0.5 and 12kbp (mean 3.7Kbp), which are the focus of this study. We have provided provisional names for these viruses following the model of Shi *et al*., (95) and the sequences have been submitted to GenBank under accession numbers MF189971-MF190055.

The majority of these virus-like contigs were related to positive sense RNA viruses (+ssRNA), including *ca*. 20 contigs from the Picornavirales, 10 Weivirus contigs, and around 5 contigs each from Hepeviruses, Nodaviruses, Sobemoviruses, and Tombusviruses. We also identified 18 putative dsRNA virus contigs (Narnaviruses, Partitiviruses and a Picobirnavirus) and 11 negative sense RNA virus (- ssRNA) contigs (5 bunya-like virus contigs, 3 chuvirus-like contigs, and two contigs each from Rhabdoviridae and Orthomyxoviridae). Our curated viruses included five DNA virus-like contigs, all of which were related to the single-stranded DNA Parvoviridae. Sequences very similar to our Caledonia Starfish parvo-like viruses 1, 2 and 3 are detectable in the publicly-available transcriptomes of *Asterias* starfish species **Fig S1;**(96). Although some of the virus-like contigs are likely to be near-complete genomes, including several +ssRNA viruses represented by single contigs of >9kbp, many are partial genomes representing only the RNA polymerase, which tends to be highly conserved (97). We identified virus-like contigs from all of the sampled taxa, although numbers varied from only three in the earthworm pool to around 40 in the sponge. This may represent differences in host species biology, but more likely reflects the different range of tissues sampled, and/or differences in sampling effort (S1 Text). A detailed description of each putative virus is provided in S2 Table.

After initially assigning viruses to potential taxonomic groups based on BLASTp similarity, we applied a maximum likelihood approach to protein sequences to infer the phylogenetic relationships of each virus. Many of the viruses derived from large poorly-studied clades recently identified by metagenomic sequencing (95,98), and most are related to viruses from other invertebrates. For example, five of the sponge picornavirales were distributed across the ‘Aquatic picorna-like viruses’ clade of Shi *et al*., (95) with closest known relatives that infect marine Lophotrochozoa and Crustacea. Associated with the breadcrumb sponge we identified sequences related to the recently described ‘Weivirus’ clade known from marine molluscs (95), and from the beadlet anemone we identified sequences related to chuviruses of arthropods (95,98). Some of the virus-like sequences were closely-related to well-studied viruses, for example Millport beadlet anemone dicistro-like virus 1 and Caledonia beadlet anemone dicistro-like virus 2 are both very closely related to Drosophila C virus (99,100) and Cricket Paralysis virus (101). Others are notable because they lack very close relatives, or because they fall closest to lineages not previously known to infect invertebrates. These include the Caledonia dog whelk rhabdo-like virus 2 sequence, which is represented by a nucleoprotein that falls between the Rabies/Lyssaviruses and other rhabdoviruses, and Barns Ness dog whelk orthomyxo-like virus 1—for which the PB2 polymerase subunit falls between Infectious Salmon Anaemia virus and the Influenza/Thogoto virus clade (Fig 2; the PA polymerase subunit shows similarity to the Thogoto viruses, but not other Orthomyxoviruses). Phylogenetic trees are presented with support values and GenBank sequence identifiers in S2 Fig, and the alignments used for phylogenetic inference and newick-format trees with support values are provided in S3 Data and S4 Data respectively.

**Fig 2.**
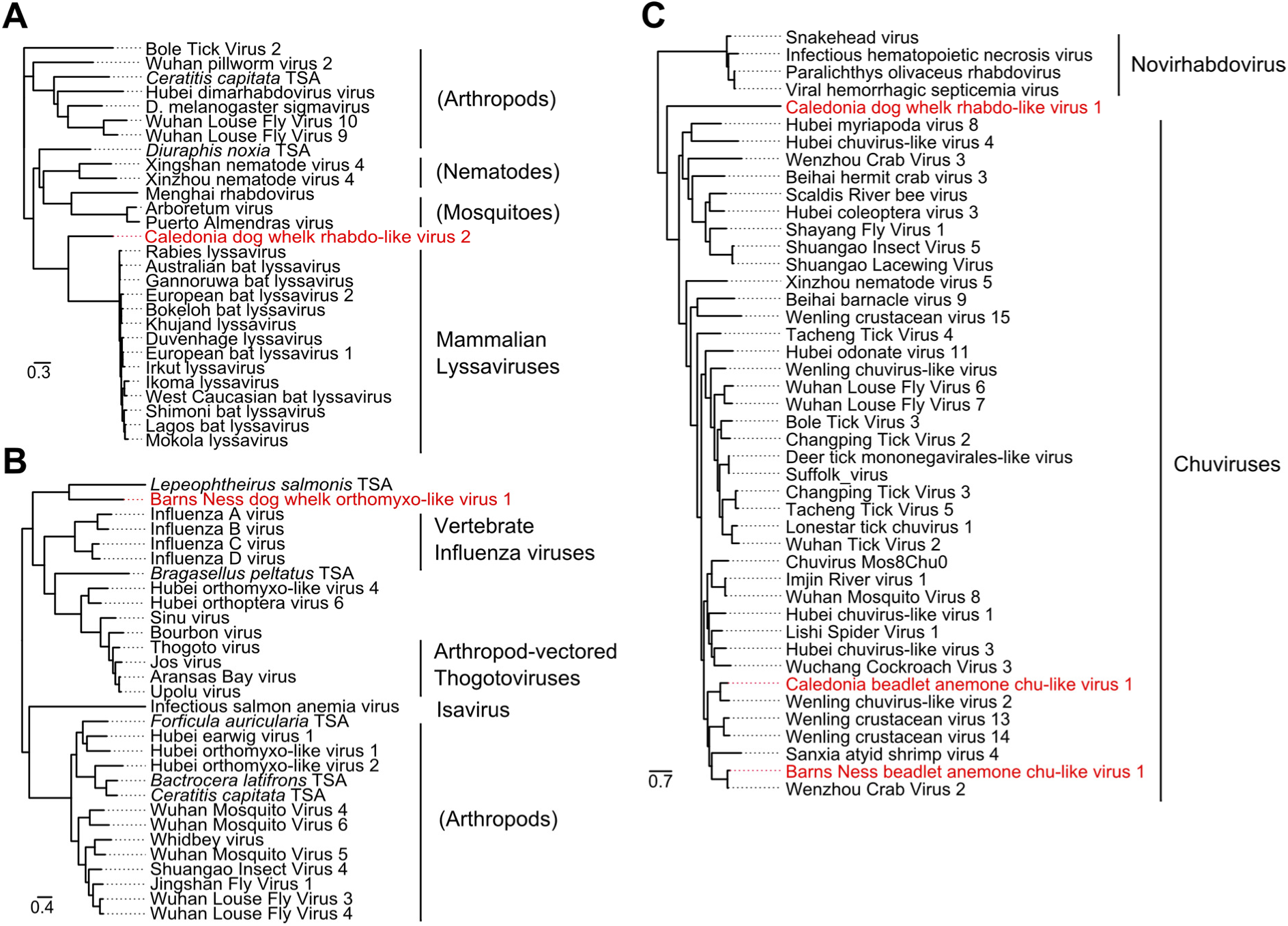
Phylogenetic relationships of virus-like contigs from the dog whelk. Mid-point rooted maximum likelihood phylogenetic trees for each of the virus-like contigs associated with viRNAs in the dog whelk (*Nucella lapillus*). New virus-like contigs described here are marked in red, sequences marked ‘TSA’ are derived from public transcriptome assemblies of the species named, and the scale is given in amino acid substitutions per site. Panels are: (A) rhabdoviruses related to lyssaviruses, inferred using the protein sequence of the nucleoprotein (the only open reading frame available from this contig, which is likely an EVE); (B) orthomyxoviruses related to influenza and thogoto viruses, inferred using the protein sequence of PB1; (C) rhabdoviruses and chuviruses, inferred from the RNA polymerase. Support values and accession identifiers are presented in S2 Fig and S4 Data, and alignments in S3 Data. Given the high level of divergence, alignments and inferred trees should be treated as tentative.

### Evidence supporting the viruses as bone fide infectious agents of the target hosts

In addition to avoiding gut content and/or nematode contamination, we sought to provide four lines of corroborating evidence that these virus-like sequences represent infections of the targeted hosts. First, we estimated the representation of potential hosts in each pool by mapping RNA-seq forward reads to the contigs of Cytochrome Oxidase 1 (COI, a highly expressed eukaryotic gene) that could be identified in our assemblies. COI reads that could not be matched to the target host species amounted to less than 0.2% of the target’s own COI reads in every case, arguing against substantial contamination with non-target taxa such as parasites or commensals. Contamination was higher in the brown alga, perhaps reflecting the challenge of recovering RNA from this taxon (S1 Text). In this case we identified around 10 contaminating taxa, amounting to 5% of the COI reads, including taxa that we might expect to live as ectocommensals on seaweeds, such as a bryozoan with 3.6% and a tunicate with 1.2%. We also identified some cross-contamination and/or adapter-switching between libraries that shared an Illumina lane (e.g. 102,103), with a mean of < 0.2% of COI reads deriving from the other libraries in the lane. Nevertheless, an average of 99.78% of the mapped COI reads in each invertebrate library derived from the targeted species (93% in the brown alga), suggesting that any viruses of contaminating species would need to be at a very high titre to be detected and erroneously attributed to the target host (read counts are provided in S3 Table).

Second, we remapped reads to the 85 focal virus contigs to measure the number of virus-derived reads relative to host COI. We reasoned that sequence reads from genuine infections are likely to appear in a single host species and to have high representation, whereas viruses present only as surface or sea-water contaminants would be present at low titre and seen in association with the multiple hosts that were collected together. We only identified one virus present at an appreciable titre in more than one host pool, suggesting that our virus-like sequences do not in general represent biological or experimental contaminants, and that the majority of viruses infected only one of the sampled host species. The exception was a 1.3 kbp partiti-like virus contig (Caledonia partiti-like virus 1), which displayed substantial numbers of reads in both the anemone and the sponge—perhaps indicative of closely related viruses infecting these highly divergent taxa. Four viruses were present at a very high level (>1% of COI in at least one library), including Caledonia beadlet anemone dicistro-like virus 2, Millport beadlet anemone dicistro-like virus, Lothian earthworm picorna-like virus 1, and in the brown alga, Barns Ness serrated wrack bunya/phlebo-like virus 1. In total, 18 of the 85 virus contigs were present at >0.1% of host COI in at least one library, and all but 8 were present at >0.01% of COI (S3 Table; S3 Fig).

Third, we recorded which strand each RNA sequencing read derived from, as actively replicating DNA viruses and -ssRNA viruses generate substantial numbers of positive sense mRNAs. As expected, all of the -ssRNA viruses in our sample (Orthomyxoviridae, Rhabdoviridae, Bunyaviridae/Arenaviridae-like, chuvirus-like) displayed substantial numbers of reads from both strands, consistent with active replication. We also detected negative-sense reads for many of the +ssRNA viruses, but not at a substantially higher rate than seen for host mRNAs such as COI (S3 Table). Nevertheless, it should be noted that although +ssRNA viruses also produce complementary (negative sense) RNA during replication, the positive to negative strand ratio is usually very high (e.g. 50:1 to 1000:1 in Drosophila C Virus), potentially making the negative strand hard to detect by metagenomic sequencing. These data provide strong evidence that all of the negative sense RNA viruses we detected comprise active infections, and are consistent with replication by the other viruses. Surprisingly, only one of the five DNA viruses (Millport starfish parvo-like virus 1) showed the strong positive sense bias expected of mRNAs, whereas the other four displayed a large negative sense bias. This suggests that these parvovirus-like sequences derived from expressed Endogenous Viral Elements (EVEs’; 104) rather than active viral infections.

Fourth, we selected 53 of the putative virus contigs for further verification by PCR (Materials and Methods; S2 Table). For most of these, we confirmed that the template was detectable by RT-PCR but not by (RT-negative) PCR, confirming that the viruses were not present in DNA form, i.e. were not EVEs (Materials and Methods; S2 Table). The exceptions were Caledonia dog whelk rhabdo-like virus 2 and (as expected) the DNA parvovirus-like contigs, which did appear in RT-negative PCR. We then estimated virus prevalence in the wild, using RT-PCR to survey all of our samples in pools of between 7 and 30 individuals. The majority of viruses had an estimated prevalence in the range 0.79-100% (S4 Table), with some virus-like sequences present in all sub-pools of the species. These ‘ubiquitous’ sequences included Caledonia dog whelk rhabdo-like virus 2, Caledonia starfish parvo-like virus 2, Caledonia starfish parvo-like virus 3, Caledonia beadlet anemone parvo-like virus 1, and 13 of the sponge viruses. This suggests that these sequences are common or that they are ‘fixed’ in the population, which could be consistent with integration into the host genome (i.e. an EVE). However, given the sampling scheme, a sponge virus at >36% prevalence has a >95% chance of being indistinguishable from ubiquitous. In addition, with the exception of Caledonia dog whelk rhabdo-like virus 2, none of the RNA viruses could be amplified from a DNA template. Taken together, the use of tissue dissection in RNA preparation, the distribution of viruses across sequencing pools, the host distribution of related viruses, the abundance and strand specificity of virus reads, the absence of DNA copies (for all but one of the RNA viruses), and the variable prevalence of the putative viruses in wild populations, support the majority of these sequences as *bone fide* active viral infections of the sampled species.

### Virus and TE-derived 21nt 5’U RNAs are present in a brown alga

Virus and TE--derived small RNAs have been well characterised in plants, fungi, and some animals, but other major eukaryotic lineages such as Heterokonta, Alveolata, Excavata and Amoebozoa have received less attention. In principle, a metagenomic approach could also be applied to these lineages, but the difficulty of collecting large numbers of individuals of a single lineage makes this challenging for single-celled organisms. Here we have taken advantage of multicellularity in the brown algae (Phaeophyceae, Heterokonta) to test for the presence of viRNAs using the serrated wrack, *Fucus serratus*. Based on a single pooled sample of tissue from 100 individuals, we identified large numbers of small RNAs with a tight distribution between 22 and 23nt, peaking sharply at 21nt. Almost all of the 21nt sRNAs were 5’ U (S4 Fig), as has been seen for sRNAs in diatoms (Bacillariophyceae, Heterokonta; 105) and is seen for some small RNA classes in green plants (106,107). Although miRNAs have been described for two other brown algae, *Ectocarpus siliculosus* (108,109) and *Saccharina japonica* (110), we were unable to identify homologues of known miRbase miRNAs among these reads. This may reflect a lack of sensitivity, as the miRNA complements of the studied brown algae are highly divergent (110), and miRNAs of *Fucus serratus* may be sufficiently divergent again to be undetectable based on sequence similarity. In contrast, 1.8% of small RNAs corresponded to the subset of high-confidence TE contigs. These small RNAs were derived from both strands, but as expected given the absence of Piwi, displayed no evidence of ‘ping-pong’ amplification—with sRNAs from both strands showing a 5’ U bias. Most interestingly, we also detected viRNAs corresponding to a -ssRNA bunya-like virus (Barns Ness serrated wrack bunya/phelobo-like virus 1; Fig 3E; S5 Fig). Although numbers were relatively small, comprising 0.01% of all small RNA reads, these were derived from both strands along the full length of the virus-like contig, peaked sharply at 21nt, and were almost exclusively 5’U. We did not detect a viRNA signature from a further two -ssRNA or from four dsRNA virus-like contigs, although their titre was very low compared to Barns Ness serrated wrack bunya/phelobo-like virus 1 (S3 Fig, S3 Table).

**Fig 3.**
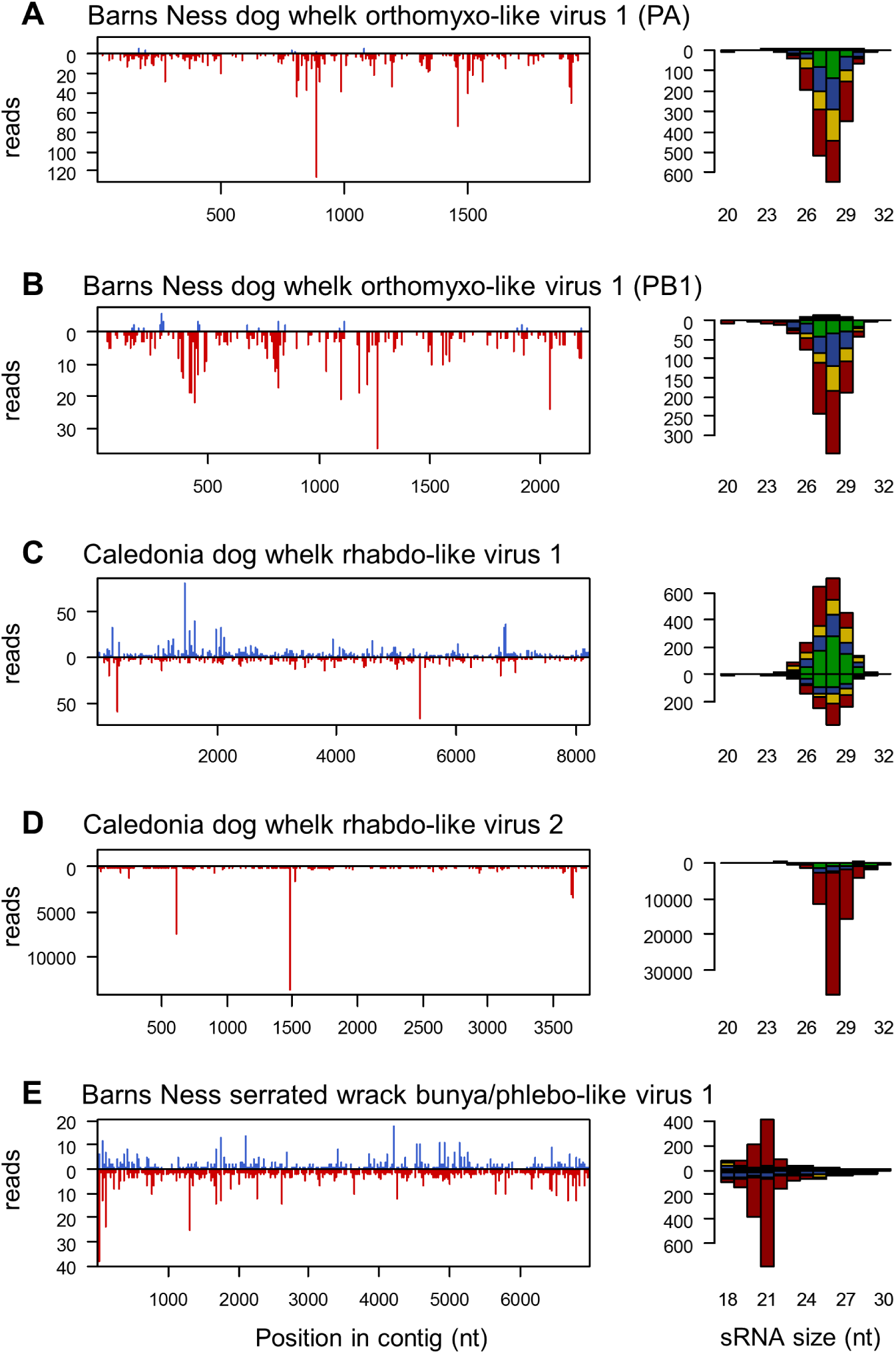
small RNAs from RNA virus-like contigs. Panels to the left show the distribution of 20-30nt small RNAs along the length of the virus-like contig, and panels to the right show the size distribution small RNA reads coloured by the 5’ base (U red, G yellow, C blue, A green). Read counts above the x-axis represent reads mapping to the positive sense (coding) sequence, and counts below the x-axis represent reads mapping to the complementary sequence. For the dog whelk (A-D), only reads from the oxidised library are shown. Other dog whelk libraries display similar distributions and the small-RNA ‘hotspot’ pattern along the contig is highly repeatable (S5 Fig). Small RNAs from the two segments of the orthomyxovirus (A and B) show strong strand bias to the negative strand and no 5’ base composition bias. Those from the first rhabdo-like virus (C) display little strand bias and no base composition bias, and those from the second rhabdo virus-like contig, which is a probable EVE (D), derive only from the negative strand and display a very strong 5’ U bias. There were insufficient reads from the positive strand of this virus to detect a ping-pong signature. Small RNAs from the four dog whelk contigs all display 28nt peaks. Small RNAs from the *Fucus* bunya/phlebo-like virus identified in the brown alga (E) derive from both strands, and show a strong 5’ U bias with a peak size of 21nt. The data required to plot the size distributions is provided in S5 Table.

### Virus-derived small RNAs are detectable in a dog whelk, but not other animal samples

Based on our knowledge of antiviral RNA interference in arthropods and nematodes we expected viral infections in our animal samples to be associated with large numbers of Dicer-generated viRNAs, with a narrow size distribution peaking between 20nt (as seen in Lepidoptera; 111) and 22nt (as seen in chelicerates, hymenopterans, and nematodes; 40,112,113). However, because animal piRNAs and viRNAs are generally modified by the addition of a 3’ 2-O-methyl group, and some nematode small RNAs are generated by direct syntehses (resulting in a 5’ triphosphate group) we additionally sequenced small RNAs treated with 5’ polyphosphatase (to remove 5’ triphosphates) and oxidised RNA (to increase the representation of small RNAs bearing a 3’ 2-O-methyl group). Furthermore, to ensure that we did not exclude viRNAs that had been edited (e.g. by ADAR; see 114), or that contained untemplated bases (e.g. 3’ adenylation or uridylation; 115), our mapping approach permitted at least two high base-quality mismatches within a 21nt sRNA. We also confirmed that remapping with local alignment, which permits any number of contiguous mismatches at either end of the read, did not substantially alter our results.

We successfully recovered abundant miRNAs in all of the animal samples, with between 20% (sponge) and 80% (starfish) of 20-23nt RNAs from untreated libraries mapping to known miRbase miRNAs (116). Consistent with the absence of a 3’ 2-O-methyl group, these miRNA-like reads had much lower representation in the oxidised libraries, there comprising only 0.4% (earthworms) to 14% (dog whelk) of 20-23nt RNAs. We also identified characteristic peaks of small RNAs derived from ribosomal RNA at 12nt and 18nt in the sponge, at 12nt and 16nt in the sea anemone, and in oxidised libraries from all organisms. The only exception to this overall pattern was for the sea anemone, in which oxidation had no effect on the number of miRNAs, although did strongly affect the overall size distribution of rRNA-derived sRNAs. This suggests the presence of a 3’ 2-O-methyl group in sponge miRNAs (S4 Fig).

However, despite our identification of more than 40 RNA virus-like contigs associated with the sponge, 17 in the sea anemone, and three in the earthworms, we were unable to detect a signature of abundant viRNAs in any of these three organisms. On average, less than 0.002% of 17-35nt RNAs from these organisms mapped to the RNA virus contigs, and those that did map were enriched for shorter lengths (17-19nt), lacked a clearly defined size distribution, and were less common in the oxidised than non-oxidised libraries (S4 Fig; S3 Table)—features consistent with non-specific degradation products, rather than viRNAs. (Note that the starfish sample lacked detectable RNA viruses, precluding the identification of RNA-virus viRNAs).

The only metazoan sample to display a clear viRNA signature was the dog whelk (*Nucella lapillus*), with 0.14% of oxidised small RNAs derived from four of the seven RNA virus-like contigs. These included both contigs of Barns Ness dog whelk orthomyxo-like virus 1, Caledonia dog whelk rhabdo-like virus 1, and Caledonia dog whelk rhabdo-like virus 2. A Narnavirus-like contig and a very low titre Bunyavirus-like contig were not major sources of viRNAs. Given the absence of detectable viRNAs in the Sponge, Sea Anemone, and Earthworm, it is notable that the viRNA-producing viruses in the dog whelk were present at a much lower copy number than many viRNA-free viruses in those organisms (e.g. Lothians earthworm picorna-like virus 1, Barns Ness breadcrumb sponge hepe-like virus 1; S3 Fig). This suggests that, had viRNAs been present in those taxa, we were likely (for many viruses) to have been be able to detect them.

Nevertheless, the virus-derived small RNAs seen in the dog whelk did not show the expected size, strand, or 2nt overhang signature of canonical Dicer-generated viRNAs (Fig 3; S5 Fig). Instead, viRNA lengths formed a broad distribution from 26 to 30nt (peaking at 28nt), more consistent with piRNAs seen in the *Drosophila* and mammalian germlines. These small RNAs were derived almost entirely from the negative-sense (i.e. genomic) strand of Barns Ness dog whelk orthomyxo-like virus 1 (Figs 3A and 3B) and Caledonia dog whelk rhabdo-like virus 2 (Fig 3D), but from both stands of Caledonia dog whelk rhabdo-like virus 2 (Figs 3C and 3E). Although this size distribution is more consistent with the piRNA pathway, only those from Caledonia dog whelk rhabdo-like virus 2 (a suspected EVE, see above) displayed the strong 5’U bias expected of primary piRNAs (Fig 3D), and none showed any evidence of ping-pong amplification. In all three cases, the putative dog whelk viRNAs were derived from the whole length of the viral genome—albeit with strong hotspots in Caledonia dog whelk rhabdo-like virus 2. Relative to miRNAs, these RNA-virus derived viRNAs were much more strongly represented in the oxidised library than the untreated library, with the miRNA:viRNA ratio increasing 300-fold—consistent with the presence of a 3’ 2-O-methyl group (S4 Fig, S5 Fig).

### The sea anemone and starfish display 5’U 26-30nt RNAs from DNA virus-like contigs

DNA viruses are a source of Dicer-mediated viRNAs in arthropods and in plants, and antiviral RNAi pathways are important for antiviral immunity to DNA viruses in both groups (reviewed in 117,118). Although our RNA sequencing strategy was intended to detect RNA viruses, we also identified four novel parvo/densovirus-like contigs (Parvoviridae; single-stranded DNA) in the starfish, and one in the sea anemone. These sequences were a substantial source of small RNAs in both organisms, particularly the starfish—contributing 0.3% of small RNAs in the untreated libraries and 3.4% of small RNAs in the oxidised library. In four of the five cases these small RNAs were almost exclusively negative sense, were 26 to 30nt in length (peaking at 28nt), and were very strongly biased toward U in the 5’ position— resembling primary piRNAs (Fig 4). However, the high prevalence and/or negative strand RNAseq bias of these four source contigs is consistent with expressed genomic integrations (EVEs) rather than active viral infections. In the other case, Millport starfish parvo-like virus 1, both positive and negative sense reads were detectable, the negative sense reads again displayed a strong 5’ U bias, but the positive sense reads displayed a postion-10 ‘A’ ping-pong signature (Fig 4B), as expected of piRNAs. Relative to miRNAs, these putative piRNAs were much more strongly represented in the oxidised library than the untreated library, consistent with the presence of a 3’ 2-O-methyl group (S4 Fig; S5 Fig).

**Fig. 4.**
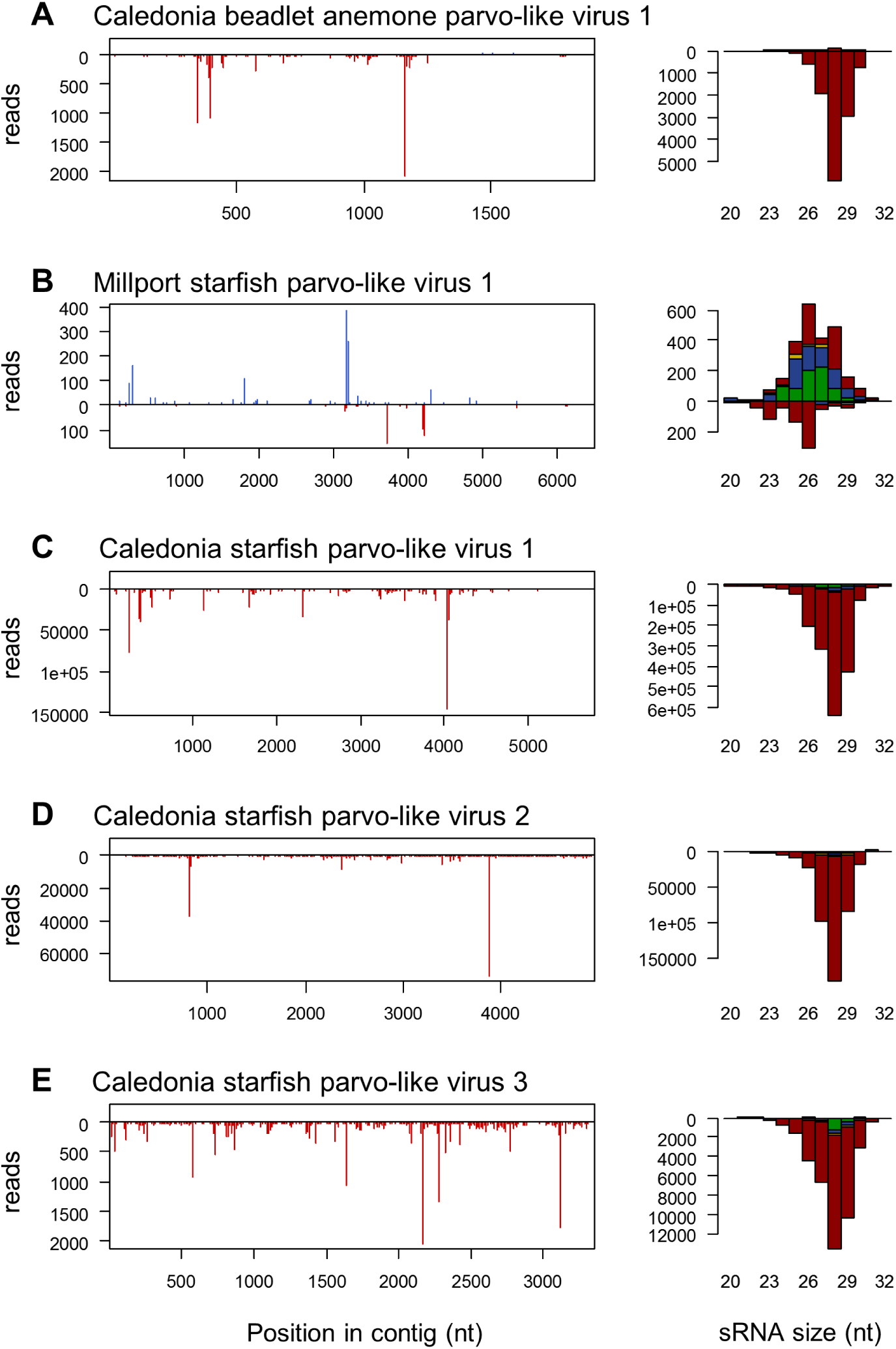
small RNAs from DNA parvo/densovirus-like contigs. Panels to the left show the distribution of 20-30nt small RNAs along the length of the parvo/densovirus-like contigs, and panels to the right show the size distribution small RNA reads coloured by the 5’ base (U red, G yellow, C blue, A green). Read counts above the x-axis represent reads mapping to the positive sense (coding) sequence, and counts below the x-axis represent reads mapping to the complementary sequence. Only reads from the oxidised library are shown, but other libraries display similar distributions, and the small-RNA ‘hotspot’ pattern is highly repeatable (S6 Fig). For all but one of the parvo/denso-like virus contigs, the small RNAs derived exclusively from the negative sense strand and showed a strong 5’U bias, consistent with piRNAs derived from endogenous copies (see main text). For one contig (B: Millport starfish parvo-like virus 1) reads derived predominantly from the positive strand and did not display a 5’ U bias. Although the number of unique small RNA sequences from this virus was small, the positive-sense small RNAs showed a slight bias to A at position 10, consistent with ping-pong (S6 Fig). The data required to plot these size distributions is provided in S5 Table.

### All of the sampled animals display somatic TE-derived piRNAs

Transposable elements and TE-derived transcripts represent a major source of piRNAs in the germlines of *Drosophila* (71), *C. elegans* (119,120), mice (121,122), and zebrafish (70), although the germline limitation seen in Drosophila has recently been shown to be derived within the Arthropods (17). Piwi-interacting RNAs are also detectable in Cnidaria and Porifera, although their tissue specificity is unclear (15), and in addition, TE transcripts in *Drosophila* and some other arthropods are also processed by Dicer to generate 21nt endo-siRNAs (17). We therefore selected a total of 146 long high-confidence TE contigs from our assemblies to analyse TE-derived small RNAs (these contigs were selected on the basis of length and similarity to repBase entries, and to best illustrate small RNA properties; contigs are provided in S5 Data). We identified large numbers of TE-derived putative piRNAs in the somatic tissues of all the sampled organisms (Fig 5). In total, between 0.17% (starfish) and 1.7% (dog whelk) of untreated small RNA reads mapped to the 146 high-confidence TE contigs (S3 Data; S5 Fig; S6 Fig). In every case except the anemone, the putative piRNAs were more highly represented in the oxidised library than in untreated or polyphosphatase-treated libraries (1.4-6% of oxidised reads), suggesting that they are 3’ 2-O-methylated and result from cleavage rather than synthesis. Despite very large numbers of piRNAs for some TE contigs, we did not observe endo-siRNA -like small RNAs similar those observed in *Drosophila* and some other arthropods (e.g. 17,123).

**Fig 5.**
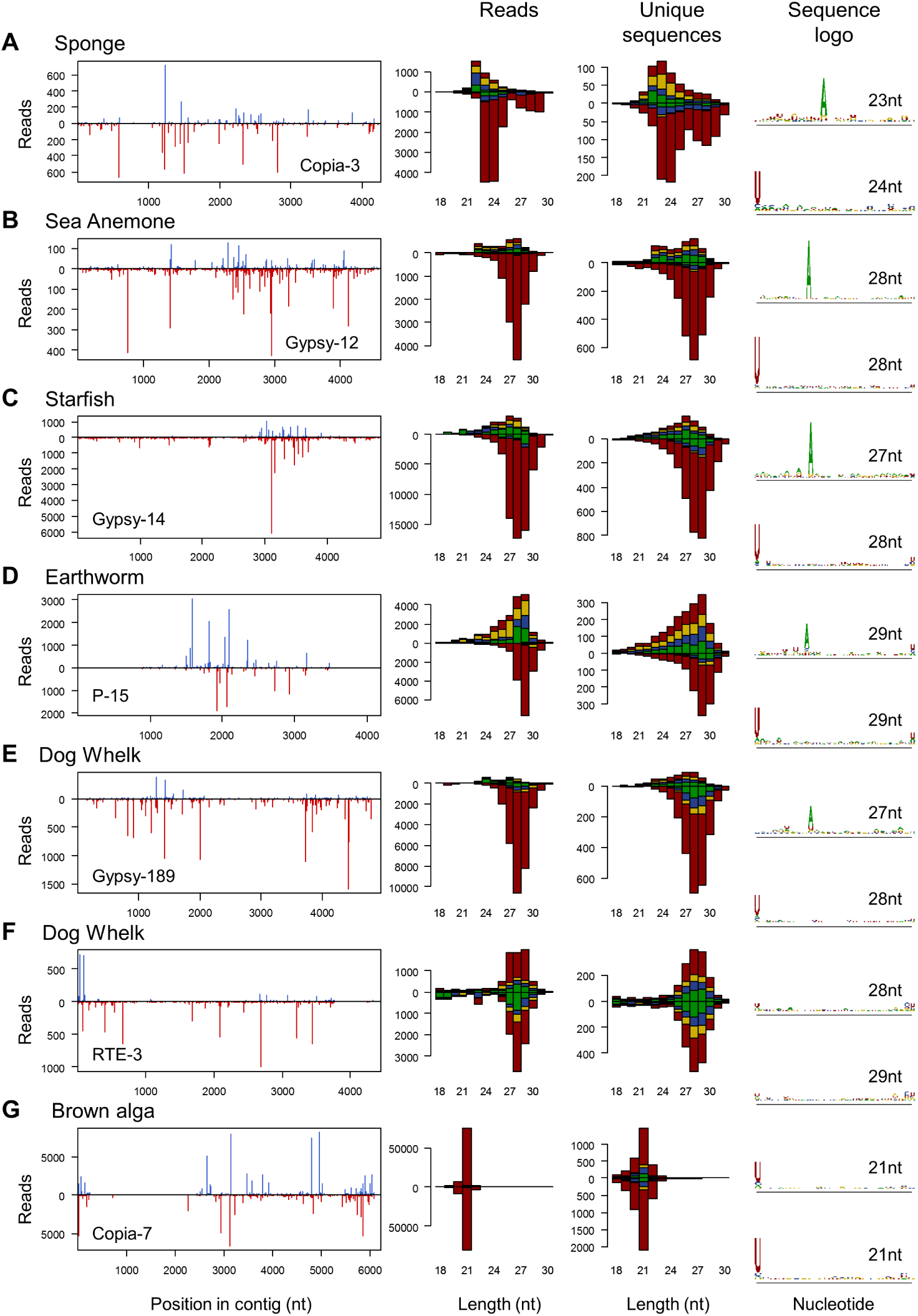
small RNAs from TE-like contigs. The four columns show (left to right): the distribution of 20-30nt small RNAs along the length of a TE-like contig; the size distribution of small RNA reads (U red, G yellow, C blue, A green); the size distribution for unique sequences; and the sequence ‘logo’ of unique sequences for the dominant sequence length. Read counts above the x-axis represent reads mapping to the positive sense (coding) sequence, and counts below the x-axis represent reads mapping to the complementary sequence. For the sequence logos, the upper and lower plots show positive and negative sense reads respectively, and the y-axis of each measures relative information content in bits. Where available, reads from the oxidised library are shown (A-F), but other libraries display similar distributions (S6 Fig). These examples were chosen to best illustrate the presence of the ‘ping pong’ signature, but other examples are shown in S6 Fig. Note that the size distribution of TE-derived small RNAs varies substantially among species, and that the dog whelk (E and F) displays at least two distinct patterns, one (F) reminiscent of that seen for some RNA virus contigs (Fig 3 C). The data required to plot these figures is provided in S5 Table.

We observed putative piRNAs derived from one or both strands of the TEs (Fig 5). Where they derived predominantly from a single strand they were generally strongly 5’U-biased (consistent with primary piRNAs). Where they derived from both strands, those from the second strand presented evidence of ‘ping pong’ amplification (i.e. no 5’ U bias, and a strong ‘A’ bias at position ten; Fig 5; S6 Fig).

However, the piRNA size distribution varied substantially among organisms and TEs. In the sponge, the length of the 5’ U-biased piRNAs either peaked at 23-24nt, or presented a broader bimodal distribution peaking at 23-24nt and 27-29nt. Where piRNAs derived from both strands, the strand with a ping-pong signature showed a shorter length distribution (22-23nt). In a few cases the putative sponge piRNAs from both strands showed a strong 5’U bias with no evidence of ping-pong amplification. In the sea anemone we consistently identified a strong peak of 5’-U biased sRNAs peaking at 28-29nt on one strand, but a generally bimodal distribution from the second ‘ping-pong’ strand (if piRNAs were present), peaking at around 23nt and 28nt. Again, both strands occasionally displayed a 5’-U bias and no evidence of ping-pong amplification. The patterns were again similar in the starfish and the earthworms, except that size distributions were unimodal, peaking at 29-30nt in the 5’-U biased strand and 25-26nt (starfish) and 26-27nt (earthworms) in the ‘ping-pong’ strand.

As with viRNAs, the only exception to this general pattern was seen in the dog whelk. In addition to TE-like contigs that displayed a classical piRNA-like signature (28nt 5’U RNAs from one strand; 26-28nt ‘ping-pong’ RNAs from the opposite strand), a small number of TE-like contigs in the dog whelk had an sRNA signature that resembled that of the dog whelk viruses Barns Ness dog whelk orthomyxo-like virus 1 and Caledonia dog whelk rhabdo-like virus 1. In these TE-like contigs, the sRNAs were derived from one or both strands, peaked broadly at 26-30nt, and lacked any bias in base composition or evidence of ‘ping-pong’ (Figs 5E and 5F). This indicates that some TEs are processed in the same way as the identified RNA viruses, (e.g. Gypsy, S6 FigD). A minority of TE-like contigs displayed an intermediate pattern, with a weak 5’U-bias from one strand, and a broad peak that lacked a pong-pong signature from the other strand. Such an intermediate pattern could result either from a single TE targeted by two different mechanisms, or from cross-mapping of sRNAs derived from different copies of the same TE inserted in different locations/contexts. As before, our permissive mapping approach and re-mapping using local alignments reduces the possibility that a large category of sRNAs escaped detection.

### The phylogenetic distribution and expression of RNAi-pathway genes

We sought to examine whether the phylogenetic distribution and expression of RNAi pathway genes in our samples was consistent with the small RNAs we observed. As expected, based on the presence of abundant miRNAs and/or an antiviral pathway, and given what is known for their close relatives (15,124–130), we identified two deeply divergent Dicer transcripts in the sea anemone, and a single Dicer transcript in each of the other animal species. The single Dicers seen in the starfish, dog whelk, and earthworms were more similar to Dicer-1 from the *Drosophila* miRNA pathway than to arthropod Dicer-2-like genes that mediate antiviral RNAi. Similarly consistent with an antiviral RNAi and/or a miRNA pathway, and with what is known for their close relatives (40,127,128,131–135), we identified two deeply divergent (non-Piwi) Argonaute transcripts in the sponge and in the anemone (S6 Table), and single Argonaute transcripts in the dog whelk and in the starfish. We identified three distinct Argonaute transcripts in the mixed-earthworm species pool, although these may represent the multiple earthworm species present. The dog whelk, starfish, and earthworm Argonautes were all more closely related to arthropod Ago-1 (which binds miRNAs but rarely viRNAs) and to vertebrate Argonautes, than to insect Ago2-like genes that mediate antiviral RNAi. It is likely that these genes mediate the miRNA pathway in these organisms, although it is possible that they may also mediate the production of novel viRNAs seen in the dog whelk. We also identified a single Dicer and Argonaute in the *Fucus*, which is consistent with what has been seen in other brown algae (108–110), and with the presence of both miRNAs and viRNAs.

Host-encoded RNA-dependent RNA polymerases (RdRp) play a key role in antiviral RNAi responses in plants (136) and nematodes (137,138). However, their role in RNAi in other animals is unknown, and they have an extremely patchy distribution across the animal phylogeny, with multiple independent losses. For example, they are absent from Vertebrata and Pancrustacea, but are present in Porifera, Cnidaria, Chelicerata, Nematoda, Bivalvia, Brachiopoda, some platyhelminthes, and non-vertebrate Deuterostomia. We identified three host RdRps in the Sea Anemone, each closely related to sequences from *Exaiptasia pallida*. We also identified a single RdRp sequence in the sponge and three in the Earthworm, although these did not cluster with their closest sequenced relatives. We were unable to identify any RdRp sequences in the dog whelk or the starfish, or in the brown alga, but it remains possible that they are present and expressed at a level too low to detect.

In animals, the piRNA pathway suppresses transposable element transcripts, and is mediated by homologs of the *Drosophila* nuclease ‘Zucchini’ and the Piwi-family Argonaute proteins Ago3 and Piwi/Aub. In mammals, fish, *C. elegans* and *Drosophila*, this pathway is primarily active in the germline and its associated somatic tissues (70,71,119–122), whereas in sponges and cnidarians—which lack a segregated germline— and many other arthropods, Piwi homologs are ubiquitously expressed (17,66,139). Consistent with our finding of TE-derived piRNAs displaying a canonical ‘ping-pong’ signature, we identified single Zucchini, Ago3 and Piwi homologs in four of the five animals surveyed (S6 Table). The exception was the sea anemone, in which we could only identify a single Piwi (more similar to *Drosophila* Piwi/Aub than to Ago-3). Surprisingly, although we did not identify canonical piRNAs in the brown alga, we did identify a possible Piwi-like transcript. However, its relatively low expression and apparent similarity to Piwi genes from the Lophotrochozoa suggest it most likely derives from the contaminating bryozoan identified by COI reads (above). Finally, consistent with the altered small RNA profile associated with oxidation, we were able to identify a single homolog of the RNA methyl transferase Hen-1 in each of the animal species, but not in the brown alga. These sequences have been submitted to GenBank under accession numbers MF288049-288076.

## Discussion

### Evidence for antiviral RNAi against -ssRNA viruses in the dog whelk and brown alga

Antiviral RNAi is an important defence mechanism in plants and many fungi, and in nematodes and arthropods, where it generates large numbers of easily detectable virus-derived small RNAs in wild-type individuals. Here we identified abundant viRNAs from RNA viruses in two of the six multicellular Eukaryotes we tested: from a bunya/phlebo-like virus in a brown alga (*Fucus serratus*) and from three different RNA virus-like contigs in the dog whelk (*Nucella lapillus*). These viRNAs displayed some, but not all, of the expected properties of a canonical antiviral RNAi response. Most strikingly, the broad length distribution around 28nt and the strong strand-bias seen in the dog whelk were not consistent with Dicer processing, which is expected to generate sRNAs from both strands simultaneously and to result in a characteristic sequence length determined by the distance between the PAZ and RNaseIII domains (140). Nevertheless, the viRNAs did display a distinct size distribution, they derived from the full length of the viral sequence, and in the dog whelk they were over-represented after oxidation— implying the presence of a 3’ 2-O-methyl group (Fig 3, S5 Fig). In addition, the viRNAs from the brown alga were notable for their very strong 5’U bias in both positive and negative sense reads. This is similar to that seen for viRNAs loaded into Arabidopsis Ago1 and Ago10 (107), perhaps reflecting a loading preference for the single Argonaute of brown algae, and is also seen for other small RNAs seen in brown algae (109,110). Taken together, we believe that these distinctive viRNA properties are consistent with an active response in both the dog whelk and the brown alga, and hence the presence of an antiviral RNAi pathway in these species—even though there is also substantial divergence from canonical arthropod antiviral RNAi and possibly from the ancestral state in Metazoa (below).

We have also considered three alternative explanations for these data. First, it is possible that the result is artefactual, and that all of the virus-like reads derive from another unknown source, such as environmental contamination. However, the large number of complementary (mRNA) sequences show the -ssRNA viruses to be active, the sequences were not identified in any of the other co-collected taxa, and the COI read counts in the dog whelk show contamination rates to be low. Contamination was higher for the brown alga, but the virus would need to be at extremely high copy number in the contaminating taxon to achieve the observed 3% of brown alga COI expression. Second, it is possible that the virus-like contigs represent expressed host loci, such as EVEs. However, sequences were not detectable by PCR in the absence of reverse transcription, and in the dog whelk the low and variable population prevalence means that any putative EVE must be segregating and at very different frequencies in different samples—more consistent with an infectious agent. Moreover, in a previous analysis of insect viruses, expressed EVEs were found to be rare relative to active viral infections: zero of 20 viruses identified by metagenomic sequencing in *Drosophila* (141). Third, even if the virus-like sequences do represent real infections, it is possible that the small RNAs do not represent an active RNAi-like response. However, their distinctive size distributions, the presence of a 3’ 2-O-methyl group in the dog whelk, and near 100% 5’U in the brown alga, argue strongly that these viRNAs are the result of active biogenesis, rather than degradation.

In contrast, it seems probable that the shorter rhabdo-like virus fragment from the dog whelk (Caledonia dog whelk rhabdo-like virus 2; Fig 3D) is a host-encoded EVE. First, the only open reading frame is homologous to a nucleoprotein and we could not detect a polymerase—despite its close relationship with the nucleoprotein of Lyssaviruses (Fig 2A). Second, RNA sequencing was dominated by negative-sense reads, suggesting a lack of mRNA expression, but consistent with host-driven expression of an integrated locus. Third, the small RNAs were exclusively negative-sense and 5’U, as sometimes seen for primary piRNAs derived from EVEs in other taxa. Fourth, the sequence was ubiquitous in our population samples, consistent with fixation and thus genome integration. Fifth, we were able to PCR amplify a band from a DNA template. If this sequence is an EVE, this could represent an alternative antiviral RNAi mechanism, akin to the piRNA-generating EVEs seen in *Aedes* mosquitoes (142).

### Evidence for substantial variation in antiviral RNAi-like responses to RNA viruses

Despite the presence of more than 70 high-confidence RNA virus-like contigs, we were unable to identify an abundant or distinct population of viRNAs derived from RNA viruses in the sponge, sea anemone, or earthworm samples (the starfish sample lacked detectable RNA viruses). Whereas the -ssRNA viruses in the dog whelk produced 1-100 viRNA reads per RNAseq read (oxidised library; S7 Fig), and Barns Ness serrated wrack bunya/phelbo-like virus 1 in the brown alga produced *ca*. 0.1 viRNA reads per RNAseq read (S7 Fig), none of the other RNA viruses gave rise to ≥0.001 viRNA reads per RNAseq read. In contrast, in an equivalent analysis of *ca*. 20 RNA viruses in wild-caught *Drosophila*, all putative viruses produced viRNAs at approximately 10-1000 viRNAs per RNAseq read (141). This represents a striking difference in the processing of RNA viruses between *Drosophila* (and other ecdysozoans, including other arthropods (17,36,37,40) and nematodes (42,43,112)), and the processing of viruses by sponges (Porifera), anemones (Cnidaria), and earthworms (Annelida). Importantly, it suggests that these animal lineages either do not process RNA viruses into small RNAs in the way that plants, fungi, nematodes or insects do, or that they do so at a level that is undetectable through the bulk small RNA sequencing of wild-type organisms and viruses—as appears to be the case for mammals (29,31–33). In either case, this suggests that the antiviral RNAi mechanisms seen in arthropods and nematodes are highly derived, and may not represent the ancestral state in Metazoa.

Nevertheless, it is necessarily hard to demonstrate that RNA viruses do *not* give rise to small RNAs in these lineages: an absence of evidence does not provide strong evidence of absence. For example, it is possible that small RNAs are abundant, but were not detected. However, this is highly unlikely as we were able to detect miRNAs, piRNAs, and small rRNAs, and we would also have detected viRNAs bearing a 5’ triphosphate or 3’ 2-O-methyl group, as well as viRNAs that had been edited or extended by untemplated bases at the 5’ or 3’ end. One alternative is that all of the RNA-virus like contigs that we identified from the sea anemone, sponge, and earthworm, were inactive and/or encapsidated at the time of collection, and thus not subject to Dicer processing. However, this is unlikely for three reasons. First, it can be ruled out for eight of the nine highest titre dsRNA viruses in the sponge, as these all showed a strong positive-strand RNAseq bias, consistent with gene expression. Second, it is not supported by the two -ssRNA virus contigs in the earthworms, which also displayed positive sense mRNA reads (although the virus copy-number was extremely low, such that that we had little power to identify either positive sense RNAseq reads or viRNAs). Finally, although the small number of negative sense reads resulting from +ssRNA virus replication makes it hard to exclude the possibility that they were inactive, it would be surprising if all of the -ssRNA viruses and dsRNA viruses (including those in the dog whelk and brown alga) were active, but none of the +ssRNA viruses were.

Perhaps a more plausible alternative is that the remaining viruses express viral suppressors of RNAi (VSRs) that completely eradicate the small RNA signature, such that it is undetectable through bulk sequencing of wild-type individuals. This appears to be the case for some mammalian viruses, where viruses genetically modified to remove their VSR do indeed form a much greater source of small RNAs (29,30,32,33). However, it is not the case for the many insect and plant viruses that express well characterised VSRs (143,144), and while it could certainly be true for some of the 80 different viruses we detect, it would be surprising if it were true for all of them.

It is also possible that abundant viRNAs are characteristic of a response against -ssRNA viruses in anemones, earthworms, and sponges, but are not characteristic of the response against +ssRNA or dsRNA viruses. This could also be consistent with our failure to detect viRNAs from putative dsRNA narnaviruses in the dog whelk and brown alga, and to a putative +ssRNA nodavirus in the brown alga. If so, then an apparent absence of antiviral RNAi in the sponge, sea anemone and earthworms may really reflect differences in the composition of the RNA virus community, with a preponderance of - ssRNA viruses in the dog whelk and their absence from the sponge or anemone. However, even if - ssRNA viruses, but not +ssRNA viruses or dsRNA viruses, give rise to viRNAs in most animal lineages, then this is still in striking contrast to the antiviral RNAi response in plants, fungi, nematodes and insects (9,36,145), and again suggests that antiviral RNAi mechanisms are highly variable among eukaryotic lineages. Finally, it also remains possible that the majority of sponges, sea anemones, and annelids do possess an active antiviral RNAi mechanism that generates abundant viRNAs from RNA viruses, but that the particular species we examined here have lost the ability. It is certainly the case that RNAi mechanisms are occasionally lost, as in one clade of the yeast genus *Saccharomyces* (22,146). However, unless antiviral RNAi is lost extremely frequently in these three animal phyla—which is not the case in arthropods or plants—it is extremely unlikely that we would by chance select three lineages that have lost the mechanism while others retained it.

### Evidence for Piwi-pathway targeting of DNA viruses in the sea anemone and starfish

We identified four parvo/denso-like virus contigs in the starfish, and one in the sea anemone. All of these sequences were detected as RNAseq reads, and were associated with abundant 26-29nt piRNA-like small RNAs (Fig 4). However, RNAseq from three of the four starfish parvo/denso-like virus contigs, and the sea anemone contig, were dominated by negative sense reads. This is hard to reconcile with the normal functioning of ssDNA parvo/denso-like viruses, which replicate via a rolling circle, and may instead reflect host-driven transcription. For these four contigs, the small RNAs were also almost exclusively negative-sense and 5’U—as expected of primary piRNAs. In contrast, RNAseq and small RNAs reads from Millport starfish parvo-like virus 1 were almost exclusively positive (mRNA) sense, with the negative strand small RNAs showing a 5’U bias and positive strand sRNAs showing weak ‘ping-pong’ signature (S5 Fig). Together, these observations suggest that at least some of parvo/denso-like virus sequences represent expressed EVEs, but also that they are targeted by a piRNA pathway-related mechanism.

Unlike for RNA viruses, we were unable to test whether these sequences represent integrations into the host genome, as integrations are indistinguishable from viral genomic ssDNA by PCR, and both +ssDNA and -ssDNA sequences are usually encapsidated by densoviruses. However, Caledonia starfish parvo-like viruses 1, 2 and 3 are nearly identical to published starfish transcripts, and the two published sequences most similar to Caledonia beadlet anemone parvo-like virus 1 are from an anemone transcriptome and an anemone genome (S2 Fig). In addition, three of the five contigs (two in the starfish, and one in the anemone) appear to be ubiquitous in our wild sample. This ubiquitous distribution and close relationship to published sequences support the suggestion (above) that some of these sequences may be host integrations. The exceptions are Caledonia starfish parvo-like virus 1 and Millport starfish parvo-like virus 1, which both had an estimated prevalence of between 4% and 20% in the larger Millport collection. We were able to recover putatively near-complete genomes of 6.5 and 5.8 Kb, containing the full length structural (VP1) and non-structural (NS1) genes, from Millport starfish parvo-like virus 1 and Caledonia starfish parvo-like virus 1, respectively (S2 Table).

If these sequences are EVEs, as seems very likely for four of the five, then their expression and processing into piRNAs may reflect the location of integration—for example, into or near to a piRNA generating locus (147,148). In contrast, if these sequences are not host EVEs, then the high expression of negative sense transcripts and the presence of primary piRNA-like small RNAs suggests an active Piwi-pathway response targeting DNA viruses in basally-branching animals. These are not mutually exclusive, and it is tempting to speculate that such integrations could provide an active defence against incoming virus infections in basal animals, as suggested for RNA-virus integrations in *Aedes* mosquitoes (142). If so, the low-prevalence Millport starfish parvo-like virus 1 sequence, which shared 72% sequence identity with Caledonia starfish parvo-like virus 1, but displayed positive sense transcripts, positive and negative sense piRNAs, and a ‘ping-pong’ signature, is a good candidate to represent an unintegrated infectious virus lineage.

### Implications for the evolution of RNAi pathways

The absence of detectable viRNAs in the sponge, sea anemone, or earthworm samples, combined with the presence of 26-29nt (non-piwi) viRNAs in the mollusc and 21nt 5’U viRNAs in the brown alga, changes our perspective of the evolution of antiviral RNAi in multicellular eukaryotes. Previously, the abundant viRNAs present in plants, fungi, nematodes and arthropods had implied that Dicer-based antiviral RNAi was ancestral to the eukaryotes and likely to be ancestral in animals, with a recent modification (or even loss; 23,24) in the vertebrates—perhaps associated with the evolution of interferons (149). Our findings now suggest three alternative hypotheses. First, antiviral RNAi may have been absent from ancestral animals, and re-evolved on at least one occasion—giving rise to the distinctively different viRNA signatures seen in nematodes, arthropods, vertebrates, and now also a mollusc. Second, the ancestral state may have been more similar to current-day mammals, which do not produce abundant easily-detected viRNAs under natural conditions, but may still possess an antiviral RNAi response (29,31–33). In this scenario, antiviral RNAi has been maintained as a defence—possibly since the origin of the eukaryotes—but has diversified substantially to give the distinctive viRNA signatures now seen in each lineage. Third, dsRNA, +ssRNA, -ssRNA, and DNA viruses may be targeted differently by RNAi pathways in basal animals, but arthropods have recently evolved a defence that gives rise to the same viRNA signature from each class. It is not possible to distinguish among these hypotheses without broader taxonomic sampling and experimental work in key lineages. For example, analyses of the Ago-bound viRNAs of Cnidaria and Porifera could help to distinguish between the first two hypotheses, and an identification of the nucleases and Argonautes and/or Piwis required for the 26-29nt mollusc viRNAs could establish whether this response is derived from a Dicer/Ago pathway or a Zucchini/Piwi like pathway. In each case, the limited taxonomic sampling and a lack of experimental data from these non-model taxa preclude any firm conclusions, and given the alternative possibilities outlined above, our interpretations should be treated as tentative. Nevertheless, the balance of evidence strongly suggests that the well-studied canonical antiviral RNAi responses of *Drosophila* and nematodes are likely to be derived compared to the ancestral state, and that there is substantial diversity across the antiviral RNAi mechanisms of multicellular eukaryotes.

The presence of piRNAs derived from transposable elements in the soma of all of the sampled animals also demonstrates a previously under-appreciated diversity of piRNA-like mechanisms. First, it argues strongly that the predominantly germline expression of the piRNA pathway in key model animals (vertebrates, *Drosophila*, and nematodes) is a derived state, and that “ping-pong” mediated TE-suppression in the soma is likely to be common in other animal phyla, as has recently been shown for arthropods (17), and has very recently been confirmed in two other molluscs (150). Second, it suggests that the TE-derived endo-siRNAs seen in *Drosophila* and mosquitoes (62,147,151–153) are absent from most phyla, and are therefore a relatively recent innovation. Third, the diversity of piRNA profiles we see among organisms—such as the bimodal length distributions of primary piRNAs in the sponge and in “ping-pong’ piRNAs in the sea anemone—suggests substantial variation among animals in the details of piRNA biogenesis. Finally, the large numbers of primary piRNAs derived from putative endogenous copies of parvo/denso-like viruses in the starfish and sea anemone, and from the putatively endogenous rhabdo-like virus 2 in the dog whelk, suggests that the piRNA processing of endogenous virus copies may be widespread across the animals, perhaps even representing an additional ancient defence mechanism.

## Materials and Methods

### Sample collections and RNA extraction

We sampled six organisms: The breadcrumb sponge *Halichondria panacea* (Porifera: Demospongiae), the beadlet anenome *Actinia equina* (Cnidaria: Anthozoa), the common starfish *Asterias rubens* (Echinodermata: Asteroidea), the dog whelk *Nucella lapillus* (Mollusca: Gastropoda), mixed earthworm species (*Amynthas* spp. and *Lumbricus* spp.; Annelida: Oligochaeta), and the brown alga *Fucus serratus* (Heterokonta: Phaecophyceae: Fucales). Marine species were sampled from rocky shores at Barns Ness (July 2014; 56.00° N, 2.45° E), and from three sites near Millport on the island of Great Cumbrae (August 2014; 55.77° N, 4.92° E) in Scotland, UK (S1 Table, S1 Text). The terrestrial sample (mixed earthworms; *Lumbricus* spp., and *Amythas* spp.), were collected from The King’s Buildings campus, Edinburgh, UK (November 2015; 55.92° N, 3.17° E). To maximise the probability of incorporating infected hosts, we included multiple individuals for sequencing (minimum: 37 sponge colonies; maximum: 164 starfish; see S1 Table for sampling details, numbers). Marine organisms were stored separately in sea water at 4°C for up to 72 hours before dissection. After dissection, the selected tissues were immediately frozen in liquid nitrogen, pooled in groups of 5-30 individuals, and ground to a fine powder for RNA extraction under liquid nitrogen (see S1 Text for details of tissue processing). Except for the brown alga *Fucus serratus*, RNA was extracted using Trizol (Life Technologies) and DNase treated (Turbo DNA-free: Life Technologies) following manufacturer’s instructions. For *Fucus*, the extraction protocol was modified from Apt *et al*., (154). Briefly, tissue was lysed in a CTAB extraction buffer, and RNA was repeatedly (re-)extracted using chloroform/isoamyl alchohol (24:1) and phenol-chloroform (pH 4.3), and (re-)precipitated using 100% ethanol, 12M LiCl, and 3M NaOAc (pH 5.2).

### Library preparation and sequencing

To avoid potential nematode contamination, an aliquot of RNA from each small (5-30 individual) pool was reverse transcribed using M-MLV reverse transcriptase (Promega) with random hexamer primers. These were screened by PCR with nematode-specific primers and conditions as described in (155) (Forward 5’-CGCGAATRGCTCATTACAACAGC; Reverse 5’-GGCGATCAGATACCGCCC). We excluded all sample pools that tested positive for nematodes from sequencing, although they were used to infer virus prevalence (below). For each host species, RNA from the nematode-free pools were combined to give final RNA-sequencing pools in which individuals were approximately equally represented. For the sponge, sea anemone, starfish, and dog whelk this pooling was subsequently replicated, using a subset of the original small pools, resulting in sequencing pools ‘A’ and ‘B’ (S1 Table, S2 Table).

Total RNA was provided to Edinburgh Genomics (Edinburgh, UK) for paired-end sequencing using the Illumina platform. Following ribosomal RNA depletion using Ribo-Zero Gold (Illumina), TruSeq stranded total RNAseq libraries (Illumina) were prepared using standard barcodes, to be sequenced in three groups, each on a single lane. Lanes were: (i) sponge, sea anemone, starfish, and dog whelk ‘A’ libraries (HiSeq v4; 125nt paired-end reads; a *Drosophila suzukii* RNAseq library from an unrelated project was also included in this lane); (ii) sponge, sea anemone, starfish, and dog whelk ‘B’ libraries (HiSeq 4000; 150nt paired-end reads); (iii) *Fucus* and Earthworms (HiSeq 4000; 150nt paired-end reads). In total, this resulted in approximately 70M high quality read pairs (i.e. after trimming and quality control) from the sponge, 60M from the sea anemone, 70M from the starfish, 70M from the dog whelk, 130M from the earthworms, and 180M from the brown alga (S3 Table).

For small RNA sequencing, total RNA was provided to Edinburgh Genomics (Edinburgh, UK) for untreated libraries (A and B), or after treatment either with a polyphosphatase (“A: Polyphosphatase”) or with sodium periodate (“B: Oxidised”). In the first case, we used a RNA 5’ Polyphosphatase (Epicentre) treatment to convert 5’ triphosphate groups to a single phosphate. This permits the ligation of small RNAs that result from direct synthesis rather than Dicer-mediated cleavage, such as 22G-RNA sRNAs of nematodes. In the second case, we used a sodium periodate (NaIO_4_) treatment (S2 Text). Oxidation using NaIO_4_ reduces the relative ligation efficiency of animal miRNAs that lack 3′-Ribose 2′O-methylation, relative to canonical piRNAs and viRNAs. This permits identification of 3′- 2′O-methylated sRNA populations, and is expected to enrich small RNA library for canonical piRNAs and viRNAs. TruSeq stranded total RNAseq libraries (Illumina) were prepared from treated RNA by Edinburgh Genomics, and sequenced using the Illumina platform (HiSeq v4; 50nt single-end reads), with all ‘A’ libraries sequenced together in a single lane, and all ‘B’ libraries sequenced together with *Fucus* and earthworm small RNAs, across four lanes. In total, this resulted in between 46M adaptor-trimmed small RNAs for the brown alga, and 150M for the sponge (S3 Table) Raw reads from RNAseq and small RNA sequencing are available from the NCBI Sequence Read Archive under BioProject accession PRJNA394213.

### Sequence assembly and taxonomic assignment

For each organism, paired end RNAseq data were assembled *de novo* using Trinity 2.2.0 (90,91) as a paired end strand-specific library (--SS_lib_type RF), following automated trimming (--trimmomatic) and digital read normalisation (--normalize_reads). Where two RNAseq libraries (‘A’ and ‘B’) had been sequenced, these were combined for assembly. For the mixed earthworm assembly, which had a large number of reads, high complexity, and a high proportion of ribosomal sequences (18%), ribosomal sequences were identified by mapping to a preliminary build of rRNA derived from subsampled data, and excluded from the subsequent final assembly. To provide a low-resolution overview of the taxonomic diversity in each sample, we used Diamond (92) and BLASTp (156) to search the NCBI nr database using translated contigs, and MEGAN6 (93) (long reads with the weighted lowest common ancestor assignment algorithm) to provide taxonomic classification. In addition, for a more sensitive and quantitative analysis of Eukaryotic contamination, we recorded the number of Cytochrome oxidase reads for each reconstructed COI sequence present. To identify cytochrome oxidase 1 (COI) sequences, all COI DNA sequences from GenBank nt were used to search all contigs using BLASTn (156), and the resulting matches examined and manually curated before read mapping. An analogous approach was taken to identify rRNA sequences, but using rRNA from related taxa for a BLASTn search.

To identify probable virus and transposable element (TE)-like contigs, all long open reading frames from each contig were identified and concatenated to provide a ‘bait’ sequence for similarity searches using Diamond (92) and BLASTp (156). Only those contigs with an open reading frame of at least 200 codons were retained. To reduce computing time, we used a two-step search. First, a preliminary search was made using translations against a Diamond database comprising all of the virus protein sequences available in NCBI database ‘nr’ (mode ‘blastp’; e-value 0.01; maximum of one match). Second, we used the resulting (potentially virus-like) contigs to search a Diamond database that combined all virus proteins from NCBI ‘nr’, with all proteins from NCBI ‘RefSeq_protein’ (mode ‘blastp’; e-value 0.01; no maximum matches). Putatively virus-like matches from this search were retained for manual examination and curation (including assessment of coverage – see below), resulting in 85 high-confidence putative virus contigs. A similar (but single-step) approach was used to search translated sequences from Repbase (94), using an e-value of 1×10^−10^ to identify TE-like contigs.

### Virus annotation and phylogenetic analysis

Translated open reading frames from the 85 virus-like contigs were used to search the NCBI ‘RefSeq_protein’ blast database using BLASTp (156). High confidence open reading frames were manually annotated based on similarity to predicted (or known) proteins from related viruses. Where unlinked fragments could be unambiguously associated based on similarity to a related sequence or via PCR (below), they were assigned to the same virus. These contigs were provisionally named based on the collection location, host species, and virus lineage. Where available, the polymerase (or a polymerase component) from each putative virus species was selected for phylogenetic analysis. Where the polymerase was not present, sequences for phylogenetic analysis were selected to maximise the number of published virus sequences available. For the Weiviruses, bunya-like viruses, and noda-like viruses, two different proteins were used for phylogenetic inference. Published viral taxa were selected for inclusion based on high sequence similarity (identifiable by BLASTp). Translated protein sequences were aligned using T-Coffee (157) mode ‘m_coffee’ (158) combining a consensus of alignments from ClustalW (159,160), T-coffee (157), POA (161), Muscle (162), Mafft (163), DIALIGN (164), PCMA (165) and Probcons (166). Alignments were examined by eye, and regions of ambiguous alignment at either end were removed. Phylogenetic relationships were inferred by maximum-likelihood using PhyML (version 20120412; Guindon & Gascuel, 2003) with the LG substitution model, empirical amino-acid frequencies, and a four-category gamma distribution of rates with an inferred shape parameter. Searches started from a maximum parsimony tree, and used both nearest-neighbour interchange (NNI) and sub-tree prune and re-graft (SPR) algorithms, retaining the best result. Support was assessed using the Shimodaira-Hasegawa-like nonparametric version of an approximate likelihood ratio test. All trees are presented mid-point rooted.

### PCR survey for virus prevalence

To estimate virus prevalence in the five animal taxa, we used a PCR survey of the small sample pools (5-30 individuals) for 53 virus-like contigs. There was insufficient RNA to survey prevalence in the brown alga. Aliquots from each sample pool were reverse transcribed using M-MLV reverse transcriptase (Promega) with random hexamer primers, and 10-fold diluted cDNA screened by PCR with primers for virus-like contigs designed using Primer3 (168,169). To confirm that primer combinations could successfully amplify the target virus sequences, and to provide robust assays, each of four PCR assays (employing pairwise combinations of two forward and two reverse primers) were tested using combined pools of cDNA for each host, with the combination that produced the clearest amplicon band chosen as the optimal assay. We took a single successful PCR amplification to indicate the presence of virus in a pool, whereas absence was confirmed through at least 2 PCRs that produced no product. PCR primers and conditions are provided in S7 Table. Prevalence was inferred by maximum likelihood, and 2 log-likelihood intervals are reported.

### RT-negative PCR survey for EVE detection

For 47 of the putative RNA virus contigs, we used PCR to verify that the sequences were not present as DNA in our sample, i.e. were not EVEs. We performed an RT-negative PCR survey of Trizol RNA extractions (which also contained DNA) using the primers and conditions provided in S7 Table. Where amplification was successful from cDNA synthesised from a DNAse-treated extraction, but not from 1:10, 1:100, or 1:0000-fold diluted RNA samples (serial dilution was necessary as excessive RNA interfered with PCR), we inferred that template DNA was absent. The remaining six (out of a total of 53 contigs for which designed PCR assays) were putative parvo/denso-like virus contigs, and were also tested as above. All six DNA virus contigs were detectable as DNA copies.

### Origin of sequencing reads and small RNA properties

To identify the origin of RNA sequencing reads, quality trimmed forward-orientation RNAseq reads and adaptor-trimmed small-RNA reads between 17nt and 40nt in length (trimmed using cutadapt and retaining adaptor-trimmed reads only; Martin, 2011) were mapped to potential source sequences. To provide approximate counts of rRNA and miRNA reads, reads were mapped to ribosomal contigs from the target host taxa and to all mature miRNA stem-loops represented in miRbase (116), using Bowtie2 (171) with the ‘--fast’ sensitivity option and retaining only one mapping (option ‘-k 1’). To identify the number and properties of virus and TE-derived reads, the remaining unmapped reads were then mapped to the 85 curated virus-like contigs, to COI-like contigs, and to 146 selected long TE-like contigs between 2kbp and 7.5kbp from out assemblies, using the ‘--sensitive’ option and default reporting (multiple alignments, report mapping quality). For small RNA mapping, the gap-opening and extension costs were set extremely high (‘--rdg 20,20 --rfg 20,20’) to exclude maps that required an indel. The resulting read mappings were counted and analysed for the distribution of read lengths, base composition, and orientation. In an attempt to identify modified or edited small RNAs, we additionally mapped the small RNA reads to the virus-like and TE-like contigs using high sensitivity local mapping options equivalent to ‘--very-sensitive-local’ but additionally permitting a mismatch in the mapping seed region (‘-N 1’) and again preventing indels (‘--rdg 20,20 --rfg 20,20’). This did not lead to substantially different results.

## Acknowledgements

We thank Andrew Rambaut, Michael Dye, Pete Doe, and Nathan Medd for assistance with field collections; Daniel Moncrieff and the Field Studies Council Scotland for logistical support in Millport; Franklin Chow, and Susan Coelho for advice on sample preparation; Helen Gunter and Karim Gharbi from Edinburgh Genomics for advice on sequencing strategy. We thank Rob Gifford, Sam Lewis and members of the Obbard lab for discussion, and Amy Buck for discussion and for contributions to the early stages of study design. We thank Prof. Benjamin tenOever and three anonymous reviewers for their constructive criticism of an earlier draft of the manuscript.

## Author contributions

Conceived and designed the experiments: DJO FMW GNS. Collected and processed field samples: FMW DJO GNS. Performed the experiments: FMW. Analysed the data: DJO FMW. Contributed reagents/materials/analysis tools: FMW DJO. Wrote the paper: FMW DJO

## Supporting information

### Figures

**S1 Fig. Taxonomic composition of contigs**.

For each of the six organisms, the coloured bars show (on a linear scale), the proportion of all Trinity contigs assigned to each major lineage using Diamond (92) and MEGAN6 (93) with ‘long reads’.

**S2 Fig. Phylogenetic trees**.

Maximum likelihood phylogenetic trees. Support values (approximate likelihood ratio test) and NCBI accession identifiers are provided. Viruses newly identified here are highlighted in red, and unannotated virus-like sequences from publicly-available transcriptome datasets are denoted ‘TSA’. Clade names follow (95,98). Alignments are provided in S3 Data and Newick format trees in S4 Data.

**S3_Fig. Relative read counts from high-titre viruses**

The bar plot shows the relative number of RNAseq reads that mapped to each of virus contigs, as a percentage relative to the read count of host COI reads (both normalised by contig length). Both positive and negative sense reads were included, from library ‘B’ only. Viruses with less than 0.01% of the COI read count were excluded. Contigs marked in bold and italic are thought to be DNA viruses or endogenous viral elements, and contigs marked with an asterisk were surveyed by (RT-PCR). Those contigs that were a source of detectable small RNAs are marked ‘viRNA’ or ‘piRNA’ as appropriate.

**S4 Fig. Size distributions of small RNAs**

Bar-plot size distributions of all small RNAs sequences. Columns correspond to species, rows to libraries. **Panel A:** All sRNAs from each library. **B**: sRNAs mapping to ribosomal sequences. Note that in most species read abundance decreases with size, indicative of degradation products, but that distinct peaks are visible in the oxidised libraries, consistent with specific short rRNAs possessing a 3’ 2-O-methyl group. **C**: sRNAs mapping to known miRNA stem-loops from miRbase (116). The proportion of putative miRNAs decreases dramatically in all oxidised libraries except the sea anemone, suggesting that miRNAs in this species possess 3’ 2-O-methyl groups. The small number of mapped miRNA reads in the brown alga is probably a result of the under-representation of close relatives in miRbase (116). **D**: sRNAs mapping to putative RNA virus contigs. Only the dog whelk has a large and distinctive distribution of virus-derived sRNAs, and these increase in the oxidised library, suggesting that they possess 3’ 2-O-methyl groups. The small number of very short virus-derived reads in the sponge are consistent with degradation products. **E**: sRNAs mapping to DNA parvovirus-like contigs. These increase in the oxidised library, suggesting that they possess 3’ 2-O-methyl groups. **F**: sRNAs mapping to selected TE-like contigs. These vary in their size range among species (21nt in the brown alga, bimodal in the sponge, peaking at 28-29nt in the other species), and increase in the oxidised library, suggesting that they possess 3’-2-O-methyl groups. Only a small proportion of TE-like contigs were used as mapping targets, and many TE-derived small RNAs remain unmapped. **G**: Unmapped sRNAs, comprising those that derived from divergent miRNAs, unrecognised viral contigs, TEs that were excluded from panel F, and all other sources. The data required to plot these figs are provided in S5 Table.

**S5 Fig. Properties and repeatability of virus-derived small RNAs**

Panels **A-D** are dog-whelk RNA viruses, panels **E-H** starfish DNA virus-like contigs, panel **I** is the anemone DNA virus, and panel **J** is the brown alga virus (note that only one library was made for this sample). In each panel, rows (top to bottom) represent each library: Library A, polyphosphatase-treated library A, Library B, and oxidised library B. Columns (left to right) are (i) Origin of reads from each genome position (red lines above the x-axis denote reads from the positive sense strand, blue lines below the x-axis denote reads from the negative sense strand; (ii) Bar plot of frequencies of unique sequences, bars above the x-axis denote reads from the positive sense strand, those below the x-axis denote reads from the negative sense strand, colours indicate 5’ base (U red, G yellow, C blue and A green); (iii) Barplot of frequencies of reads; (iv) Sequence logo for the unique sequences of the most frequent length deriving from the positive strand; (v) Sequence logo for the unique sequences of the most frequent length deriving from the negative strand. The data required to plot the size distributions are provided in S5 Table.

**S6 Fig: Properties and repeatability of TE-derived small RNAs**

Panels **A-R** show the small RNA properties of selected high-confidence TE-like contigs: starfish panels **A-C**, dog whelk **D-F**, sponge **G-I**, earthworms **J-L**, sea anemone **M-O**, brown alga **P-R**. Rows and columns are as in S5 Fig. The data required to plot the size distributions are provided in S5 Table.

**S7 Fig. RNAseq and sRNA reads per metagenomic contig**

For each metagenomic contig (pale grey) the ratio of sRNAs (20-31nt) to RNAseq reads is shown on the x-axis, and the ratio of 20-24nt sRNAs (expected viRNAs) to 25-31nt sRNAs (expected piRNAs) is shown on the y-axis. Contigs are only included if they are >0.75Kbp in length and produced at least 20 small RNAs; Contigs in dark grey have sequence similarity to known TEs, and contigs in colour correspond to the curated viruses. Based on *Drosophila*, TEs (dark grey) are expected to appear in the lower right quadrant of each plot, and viruses (colour) in the upper right (compare with Fig 4 in 141). Only the dog whelk (panel **A**) and the brown alga (panel **D**) display sRNAs from RNA virus contigs, although DNA virus-like contigs display piRNA-like small RNAs in the sea anemone (panel **C**) and the starfish (panel **B**). No other viruses produced sufficient viRNAs to appear on these figs. All figs (except the brown alga) use data from RNAseq library B and the corresponding oxidised sRNAs (which is enriched for viRNAs over miRNAs), and sRNA counts exclude those mapping to known (miRbase) miRNAs and rRNAs.

### Tables

**S1 Table. Sample collection details**.

Detailed descriptions of the sample collection locations, dates and numbers of individuals sampled for each target taxon, along with sample pool information, including which extraction pools were included in sequencing pools, and which were excluded due to detection of suspected nematode contamination.

**S2 Table. Detailed descriptions of putative viruses and virus-like contigs**.

Detailed descriptions of the candidate virus fragments identified by protein similarity search, including phylogenetic position, estimated prevalence, approximate coverage, ORF number and most similar viral proteins identified by BLASTp, detectability by RT-negative PCR, GenBank accession numbers, and additional notes.

**S3 Table. Sources of RNAseq and small RNA reads**.

Cytochrome oxidase coverage relative to that of the target taxon, and virus coverage for the target viruses (positive and negative strand) relative to that of COI.

**S4 Table. Virus Prevalence**

Estimated virus prevalence inferred by maximum likelihood (with 2 log-likelihood intervals) from an RT-PCR survey of pooled samples (methods as in 141).

**S5 Table. Size Distribution of small RNAs**

Raw counts necessary to plot Fig 3, Fig 4, Fig 5, S4 Fig, S5 Fig, S6 Fig

**S6 Table. RNAi related genes identified from organisms**

Counts of key RNAi related genes identified in transcriptomes of target taxa along with GenBank accession numbers for sequences.

**S7 Table. PCR primers and conditions**

PCR primer names and sequences, thermocycler conditions, and PCR recipes for virus prevalence, and RT-negative (EVE detection), assays.

### Data

**S1 Data. Raw de novo-assembled contigs**

For each of the six species pools, the raw meta-transcriptomic contigs generated by Trinity are provided in compressed (gzipped) fasta format. The majority of contigs are likely to derive from the named host species and associated microbiota, although there may be a small amount of cross contamination among libraries run in the same lane. These contigs have not been curated, and are likely to include chimeric assemblies. As such, they are not suitable for submission to GenBank, and should be treated with caution.

**S2 Data. Putative virus-like contigs**

Raw meta-transcriptomic contigs generated by Trinity that have detectable sequence similarity (using Diamond) to virus proteins in GenBank, provided in compressed (gzipped) fasta format. Contig titles are annotated using the species name of the top match, followed by the percentage identity of that match in the sequence, and the e-value associated with that match. The contigs have not been curated, and are likely to include chimeric assemblies. As such, they are not suitable for submission to GenBank, and should be treated with caution.

**S3 Data. Protein sequence alignments**

Protein sequence alignments used for phylogenetic analyses are provided in compressed (gzipped) gapped fasta format, with regions of poor alignment (identified by eye) deleted. Sequence titles comprise the taxon name and NCBI accession identifier for the sequence.

**S4 Data. Phylogenetic trees**

Phylogenetic trees are provided in compressed (gzipped) newick format. Sequence titles comprise the taxon name and NCBI accession identifier for the original protein sequence.

**S5 Data. Long high-confidence TE-like contigs**

Selected meta-transcriptomic contigs generated by Trinity that have detectable sequence similarity (using Diamond) to TEs in Repbase (94), provided in compressed (gzipped) fasta format. Contig titles are annotated using the host species name and the top-match TE in Repbase.

### Text

**S1 Text. Sampling, tissue preparations and RNA extractions**

Detailed description of the sampling, tissue preparations and RNA extractions techniques employed for each target taxon

**S2 Text. RNA oxidation treatment**

Protocol for sodium periodate (NaIO4) oxidation of RNA prior to library preparation, to enrich small RNA libraries for canonical piRNAs and viRNAs by reducing the relative ligation efficiency of metazoan miRNAs that lack 3′-Ribose 2′O-methylation.

